# Topical ABT-263 treatment reduces aged skin senescence and improves subsequent wound healing

**DOI:** 10.1101/2024.08.19.608670

**Authors:** Maria Shvedova, Rex Jeya Rajkumar Samdavid Thanapaul, Joy Ha, Jannat Dhillon, Grace H Shin, Jack Crouch, Adam C Gower, Sami Gritli, Daniel S Roh

## Abstract

Senescent cells (SnC) accumulate in aging tissues, impairing their ability to undergo repair and regeneration following injury. Previous research has demonstrated that targeting tissue senescence with senolytics can enhance tissue regeneration and repair by selectively eliminating SnCs in specific aged tissues. In this study, we focused on eliminating SnC skin cells in aged mice to assess the effects on subsequent wound healing. We applied ABT-263 directly to the skin of 24-month-old mice over a 5-day period. Following topical ABT-263, aged skin demonstrated decreased gene expression of senescent markers p16 and p21, accompanied by reductions in SA-β-gal and p21-positive cells compared to DMSO controls. However, ABT-263 also triggered a temporary inflammatory response and macrophage infiltration in the skin. Bulk RNA sequencing of ABT-263-treated skin revealed prompt upregulation of genes associated with wound healing pathways, including hemostasis, inflammation, cell proliferation, angiogenesis, collagen synthesis, and extracellular matrix organization. Aged mice skin pre-treated with topical ABT-263 exhibited accelerated wound closure. In conclusion, topical ABT-263 effectively reduced several senescence markers in aged skin, thereby priming the skin for improved subsequent wound healing. This enhancement may be attributed to ABT-263-induced senolysis which in turn stimulates the expression of genes involved in extracellular matrix remodeling and wound repair pathways.

## Introduction

Accumulation of senescent cells (SnC) within multiple tissues during aging has a strong association with age-related decline and disease ^1–3^. These chronic SnCs, although relatively rare, can be generally characterized by expression of cell cycle inhibitors such as p16, p21, increased senescence-associated beta-galactosidase (SA-β-gal), and a senescence-associated secretory phenotype (SASP). The clearance of chronic SnCs is expected to decrease chronic, low-grade inflammation and improve tissue repair capacity, however, their effects can be both tissue and age specific ^4^. In aged skin, chronic SnCs are reported to comprise up to 15% of cells in the epidermis and dermis in aged humans, primates, and mice ^5-8^. The influence of these chronic SnCs residing in aged skin on cutaneous wound repair has not been well studied, but is hypothesized to contribute to the wound healing delay that occurs in aging^9^.

Senotherapeutics represent a novel therapeutic approach that has spawned from the development of agents and strategies that specifically target cellular senescence as one of the potential mechanisms of aging and age-related diseases ^1,10^. Many senolytics, drugs that eliminate SnCs, are already in human clinical trials, demonstrating early promise of ameliorating age-related diseases such as osteoporosis, kidney disease, and pulmonary fibrosis^11^. These approaches are focused on removing chronic disease-associated SnCs but do not emphasize how senescence-reduction in aged tissues might affect responses to a subsequent injury such as cutaneous wounds. Targeting chronic SnCs in aged mouse skin through in vitro and in vivo approaches have demonstrated some reduction in detectable SnCs ^12^, however, there is limited data on the impact of SnC removal on subsequent wound repair.

We specifically examined ABT-263, a potent and broad-spectrum senolytic that inhibits the anti-apoptotic mechanisms of SnCs by inhibiting the anti-apoptotic proteins BCL-2 and BCL-xL, which are upregulated in SnCs. ABT-263 has shown effectiveness in targeting aged dermal fibroblast and can alter aged skin senescence markers ^9,12^. However, the impact of ABT-263 on aged wound healing has not been thoroughly investigated.

## Results

### Topical ABT-263 reduces general senescence markers in aged mouse skin

The skin of aged (24-month-old) mice underwent a 5-day topical treatment with either 5μM ABT-263 or the control vehicle DMSO alone. The application of ABT-263 resulted in a significant decrease in gene expression of p16 and p21 in the skin compared to DMSO treatment (Figure 1A). Additionally, there was a reduction in the number of both p21 and SA-β-gal positive cells following ABT-263 treatment compared to DMSO (Figure 1B, C). Young (2-month-old) mice were treated in similar manner and did not demonstrate any significant decrease in expression of p16 or p21 (Supplemental Figure 1).

**Figure 1.**
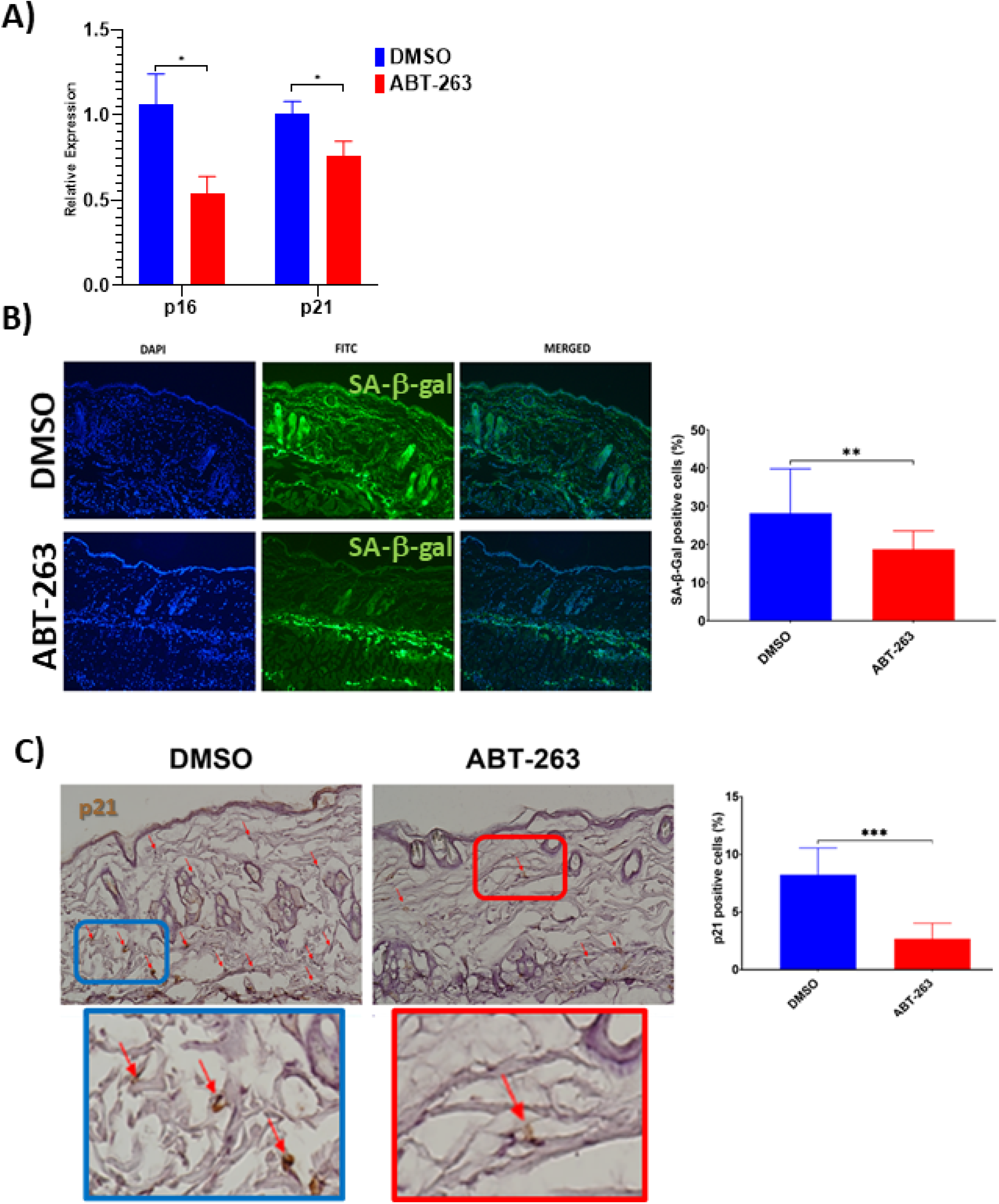
Decreased senescence markers in aged skin treated with topical ABT-263. A) relative p16 and p21 expression in whole skin after 5d of ABT-263 (N=5) vs DMSO treatment (N=5). B) SA-β-gal positive cells (green) and quantification of percentage of positive cells. C) p21 staining and quantification of percentage of positive cells. *p<0.05, **p<0.01, ***p<0.001, t-test.

### ABT-263 treatment acutely increases inflammation and dermal macrophage recruitment

There was evidence of increased cellular infiltration in both ABT-263 and DMSO-treated aged skin, with a higher level of infiltrating cells per high power field observed in the ABT-263 group (Figure 2A,B). This inflammation was driven by an elevated presence of F4/80+ macrophages after ABT-263 treatment located in both the superior papillary and lower reticular dermis (Figure 2C, D). Examination of additional immune cells demonstrated fewer CD4+ and significantly fewer CD8+ T-cells in aged skin treated with topical ABT-263 (Supplementary Figure 2).

**Figure 2.**
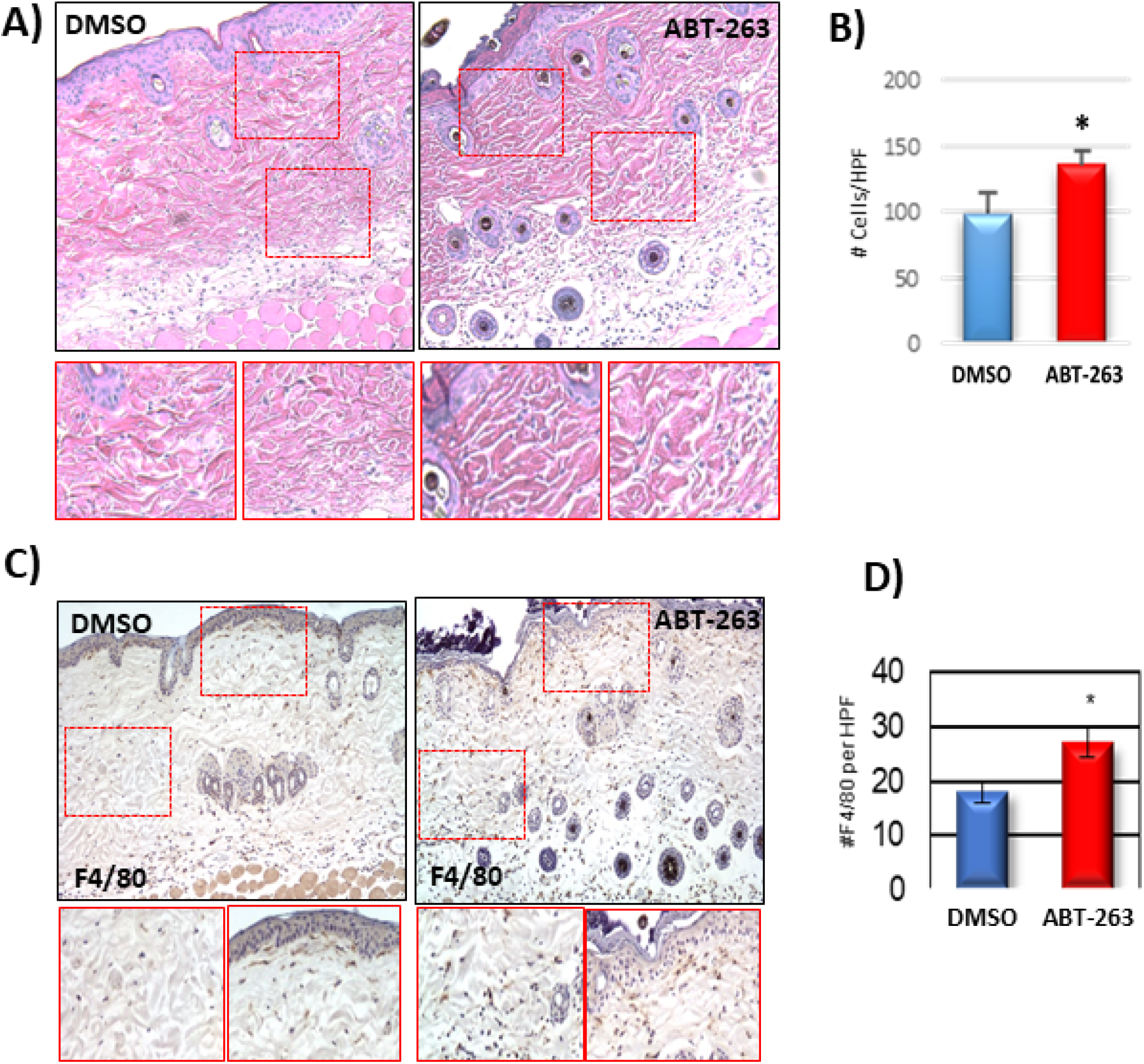
Increased dermal macrophage infiltration in aged skin treated with ABT-263. A) H&E sections of aged skin after 5d of ABT-263 (N=5) vs DMSO (N=5) treatment. B) Number (#) of total cells/high powered field (HPF). C) F4/80 staining with areas of magnification in red displayed below, and D) quantification of number (#) of F4/80+ cells per HPF. * indicates p<0.05, t-test.

### ABT-263 drives significant gene expression changes in skin including increased apoptotic signaling and mitochondrial dysfunction

Bulk RNA sequencing of ABT-263 and DMSO treated skin revealed strong and significant differential gene expression between the two groups (Supplementary Table 1, Figure 3A). The elevated expression of annexins and caspases and decreased expression of *Bcl2* indicate that ABT-263 promotes apoptosis by inhibiting anti-apoptotic Bcl-2 family proteins. We also used Gene Set Enrichment Analysis (GSEA) to measure the coordinate regulation of gene sets related to pathways and processes (Supplementary Table 2). GSEA confirmed that apoptotic and mitochondrial electron transport chain (ETC) gene sets were significantly coordinately up- and down-regulated, respectively, with respect to ABT-263 treatment (FDR *q* < 0.05, Figure 3B). A heatmap of the expression of *Bcl2* and the most significant genes within each gene set is shown in Figure 3C.

**Figure 3.**
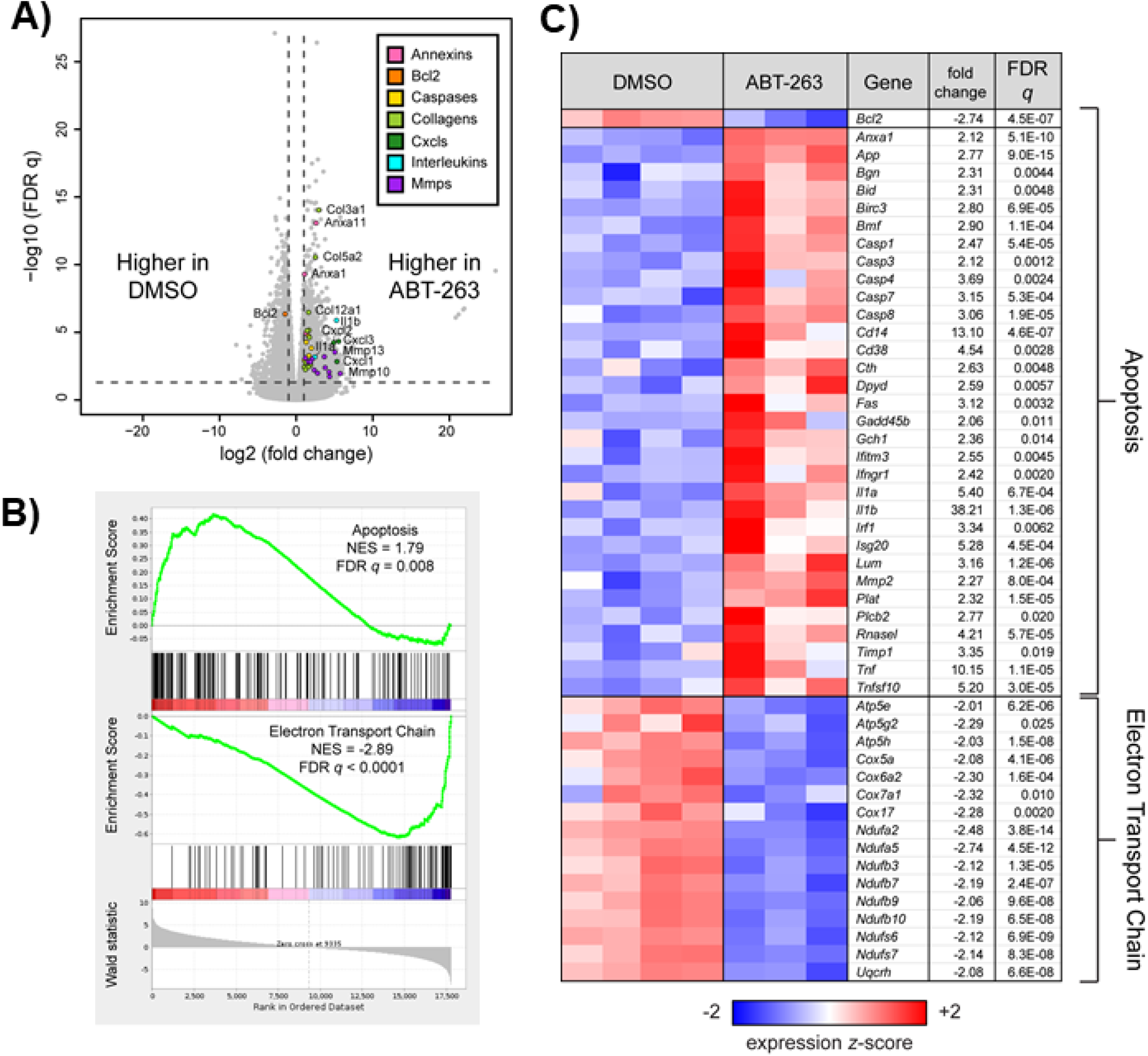
ABT-263 induced changes in aged skin apoptosis and mitochondrial oxidative phosphorylation. A) Volcano plot corresponding to a Wald test of RNA sequencing data obtained from aged skin treated with DMSO (n=4) or ABT-263 (n=3). Selected genes with significant and strong differential expression (FDR q < 0.05, |fold change| > 2, indicated by dashed lines) are colored according to their function. B) Gene Set Enrichment Analysis (GSEA) enrichment plots showing significant (FDR *q* < 0.05) coordinate up- and down-regulation, respectively, of apoptosis (Hallmark) and mitochondrial electron transport train (WikiPathways, WP295) in ABT-263-vs DMSO-treated aged skin. C) Heatmap of *Bcl2* and leading edge genes from each gene set that are also individually significant and strongly regulated by ABT-263 (Wald FDR *q* < 0.05, |fold change| > 2). Fold changes for downregulated genes are represented with their negative reciprocal (i.e., values of +2 and −2 correspond to two-fold up- or down-regulation by ABT-263, respectively). Genes are ordered alphanumerically by symbol within each gene set. Variance-stabilizing-transformed expression values are z-score-normalized to a mean of zero and SD of 1 across all samples in each column, with blue, white and red indicating z scores of ≤ 2, 0, and ≥ 2, respectively.

### ABT-263 increases the expression of many, but not all, SASP factors

To determine the effect of topical ABT-263 on SASP factors, we first used GSEA to determine whether the murine SASP gene set (R-MMU-2559582) in the Reactome pathway database was coordinately regulated with respect to treatment. The expression of these genes was not significantly skewed in either direction by ABT-263 treatment (*p* = 0.93, Figure 4A). As the Reactome SASP gene set is primarily composed of genes involved in the anaphase promoting complex and cyclin regulation, with few encoding secreted proteins, we then performed GSEA using the mouse homologs of a completely separate set of genes compiled by Coppé et al^13^ that are increased in SASP, including interleukins, chemokines, inflammatory and growth factors, proteases/antiproteases, and soluble receptors/ligands. These genes were strongly coordinately upregulated by ABT-263 treatment (*p* < 0.001, Figure 4B). A heatmap of the expression of all SASP-related genes listed in Coppé et al is shown in Figure 4C, indicating that, while most of these genes are upregulated by ABT-263, others (e.g., IGFBP family members) are unchanged or downregulated with respect to treatment.

**Figure 4.**
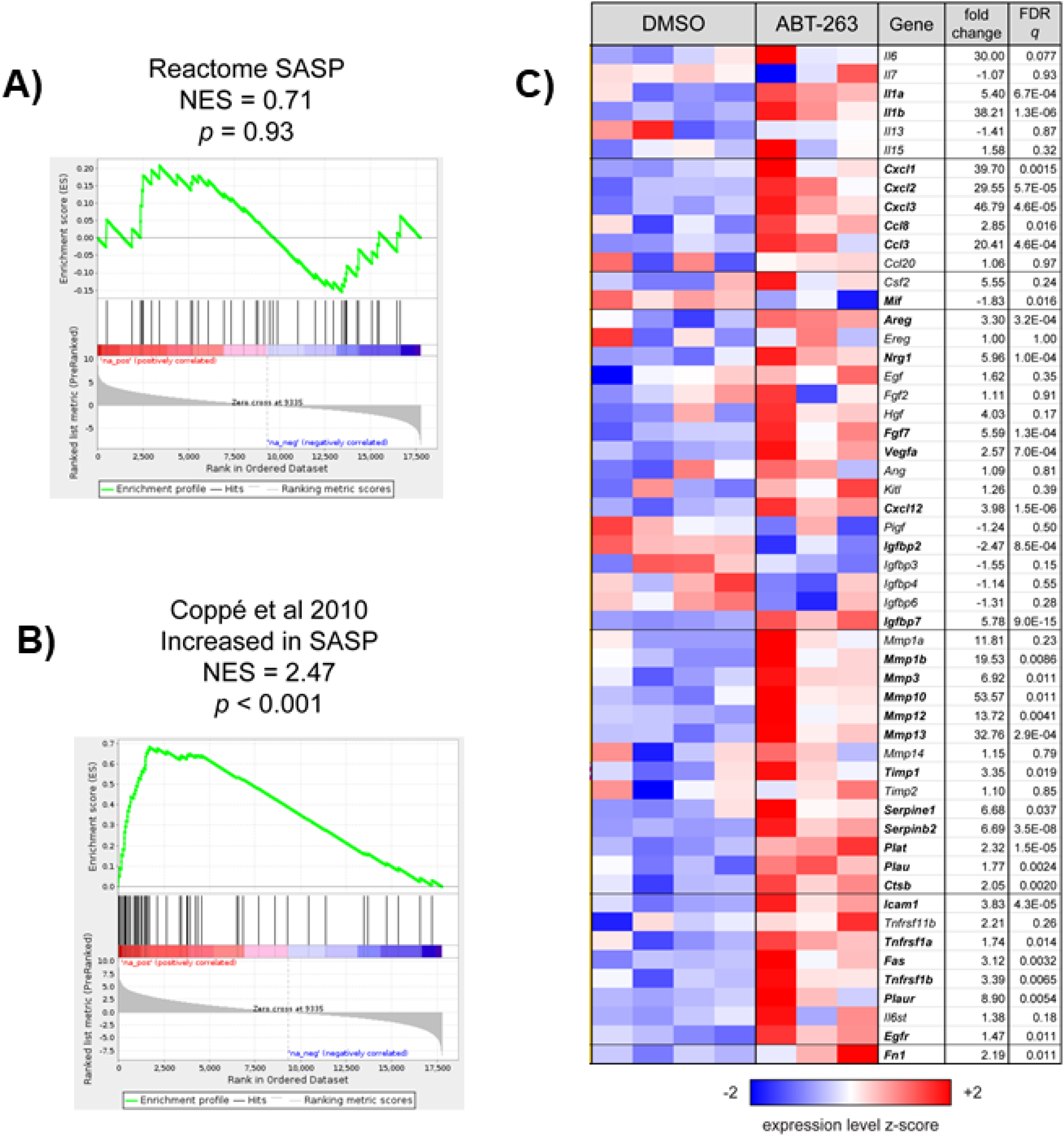
ABT-263 induced changes in the expression of some SASP factors. A) Gene Set Enrichment Analysis (GSEA) enrichment plot for Reactome SASP gene set, which was not coordinately regulated with respect to ABT-263 treatment (*p* = 0.93). B) GSEA enrichment plot showing that a set of 54 genes that are characterized as “increased” or “increased or no change” with SASP in Table 1 of Coppé et al^13^ are significantly coordinately upregulated with ABT-263 treatment (*p* < 0.001). C) Heatmap of all genes described as in Coppé et al^13^. Genes are presented in the same order and groups as in the source table; those with symbols in boldface are significantly regulated by ABT-263 (Wald FDR *q* < 0.05) in this experiment. Fold changes for downregulated genes are represented with their negative reciprocal (i.e., values of +2 and −2 correspond to two-fold up- or down-regulation by ABT-263, respectively). Variance-stabilizing-transformed expression values are z-score-normalized to a mean of zero and SD of 1 across all samples in each column, with blue, white and red indicating z scores of ≤ 2, 0, and ≥ 2, respectively.

### Topical ABT-263 upregulates multiple signaling pathways relevant to wound healing

We also used GSEA to examine the regulation of several gene sets representing the multiple stages of wound healing. In ABT-263 treated skin, genes related to hemostasis, inflammation, proliferation, angiogenesis, and collagen synthesis with extracellular matrix regulation were all coordinately upregulated (Figure 5).

**Figure 5.**
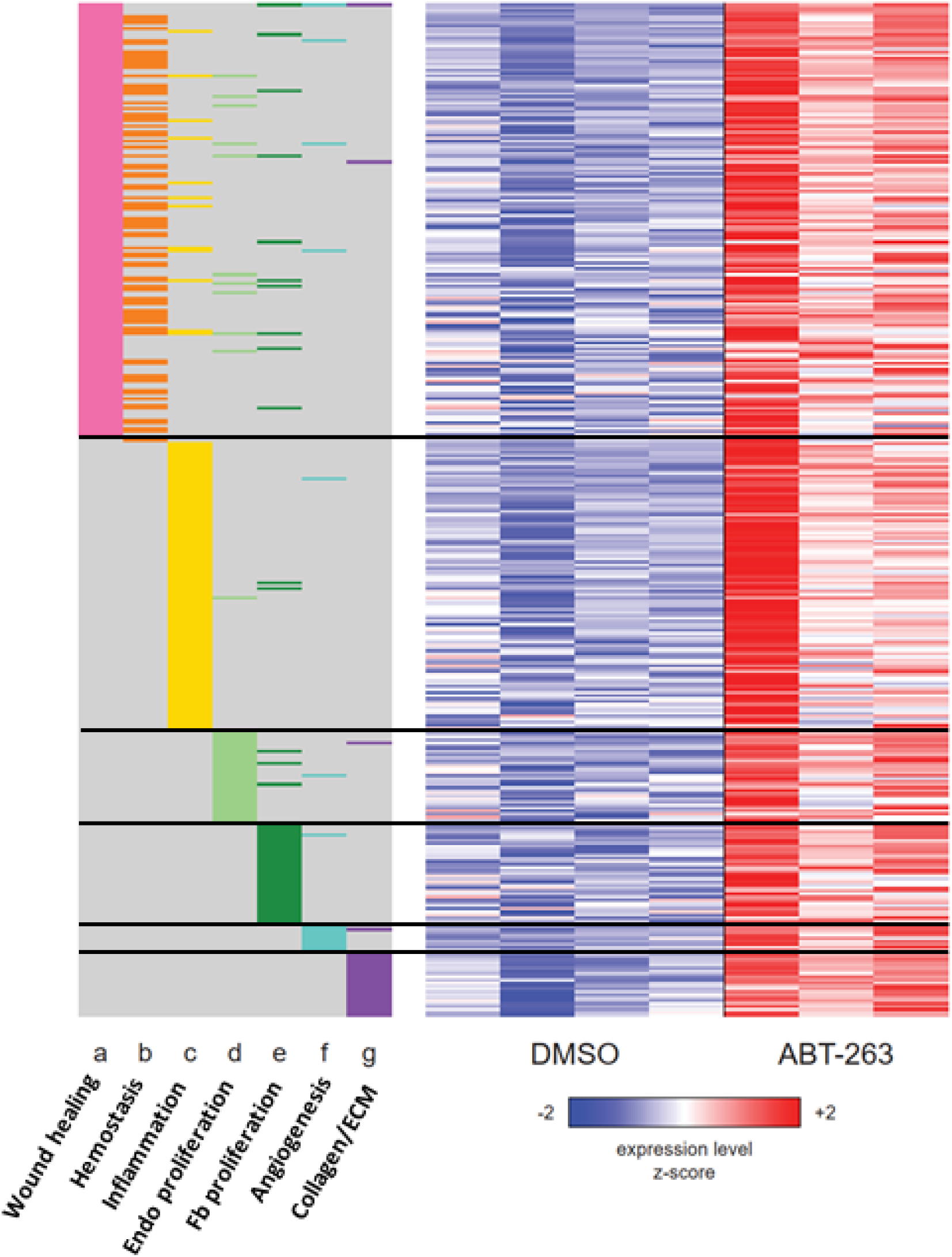
Heatmap of the union set of leading edge genes from seven wound healing related gene sets with significant coordinate upregulation (FDR *q* < 0.05) in ABT-263-vs DMSO-treated aged skin. The membership of each gene within each gene set is indicated with a colored box at the side of the heatmap: wound healing (GO:0042060), hemostasis (GO:0007599), inflammatory response (Hallmark), positive regulation of endothelial cell proliferation (GO:0001938), fibroblast proliferation (GO:0048144), angiogenesis (Hallmark), collagen biosynthesis and modifying enzymes (Reactome R-MMU-1650814). Rows and columns correspond to genes and samples, respectively. Rows are sorted from top to bottom first by gene set and then by Wald statistic. Variance-stabilizing-transformed expression values are *z*-score-normalized to a mean of zero and SD of 1 across all samples in each column, with blue, white and red indicating *z* scores of ≤ 2, 0, and ≥ 2, respectively.

### Pretreatment of aged skin before injury improves subsequent time to wound closure

Given the substantial upregulation of wound-healing-related genes and pathways following topical ABT-263 treatment, we assessed the impact of topical ABT-263 treatment on subsequent wound healing in aged mice. 24-month-old male mice received either 5μM ABT-263 in DMSO or DMSO for 5 days followed by creation of a 1cm full-thickness dorsal skin wound (Figure 6A). Mice treated with ABT-263 exhibited a significantly accelerated time to achieve complete wound closure, which became statistically significant by day 15 (Figure 6B,C). 33% of ABT-263 treated mice had completely healed by day 18, in contrast to 0% of DMSO treated mice (Figure 6D). By Day 24, 80% of ABT-263 treated mice had completely healed compared to only 56.3% in the DMSO group, representing a 1.4x fold improvement in complete healing rate.

**Figure 6.**
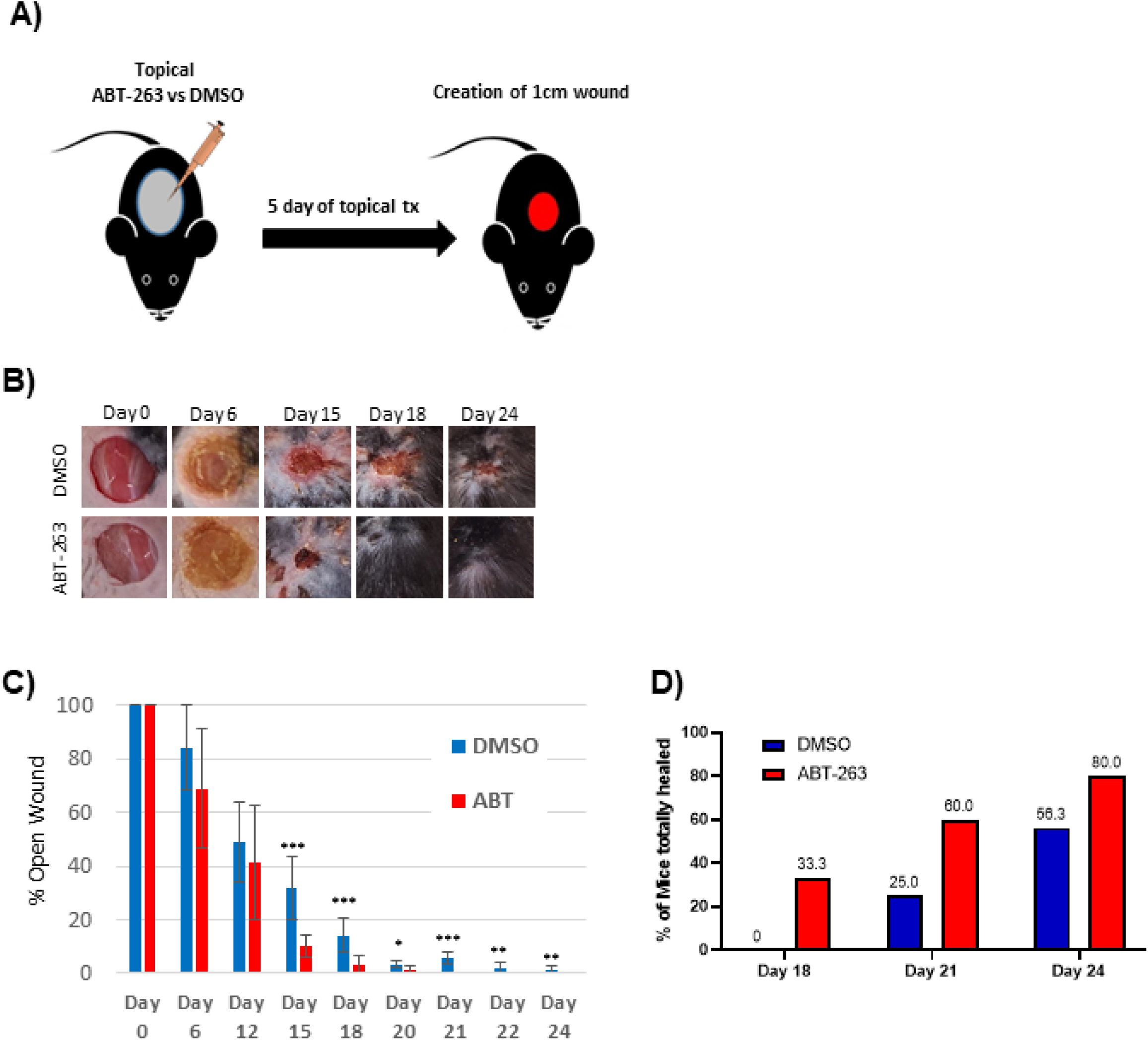
ABT-263 skin pre-treatment accelerates wound healing in aged mice. A) Schematic of the experiment. B) Representative wound photos immediately after 5d ABT-263 vs DMSO treatment. ABT-263 (N=5-8 per timepoint) vs DMSO (N=5-8 per timepoint). C) % wound contraction. D) % of aged mice with completely healed wounds. t-test, *p<0.05, **p<0.01, ***p<0.001.

## Discussion

Aged skin demonstrates cellular, molecular, and structural changes with accumulation of chronic SnCs ^1,5,10^. As a result, aged skin is characterized by decreased barrier function, attenuated dermal thickness and altered structure, and an overall impaired regenerative capacity. Delays in wound healing in aging are multifactorial; however, the intrinsic cellular changes, with accumulation of chronic SnCs, may contribute to delayed repair. Our study demonstrates that the removal of chronic skin SnCs with topical ABT-263 senolytic application can enhance regenerative capacity, leading to accelerated cutaneous wound healing.

We selected ABT-263 for this study due to its reported broad-spectrum senolytic activity on multiple cell types^12,14-18^. Furthermore, prior studies have utilized ABT-263 with robust effect on skin-specific SnC clearance of fibroblasts, myofibroblasts, and melanocytes ^12,18,19^. To limit potential systemic effects such as transient neutropenia and thrombocytopenia^20^, we utilized topical applications of ABT-263 dissolved in DMSO versus DMSO vehicle alone. After 5 days of treatment we observed no significant changes in serum platelets and actually found elevated neutrophils in ABT-263 treated aged mice compared to DMSO controls (Supplementary Figure 3). Thus, topical ABT-263 treatment appears to avoid the acute negative systemic effects of neutropenia and thrombocytopenia.

Topical application of ABT-263 modified apoptosis and mitochondrial oxidative phosphorylation in aged skin, confirming its known targeted effects on other tissues and cell types. SnCs upregulate Bcl-2 to maintain an anti-apoptotic state, which ABT-263 targets to activate the apoptotic signaling pathway and remove SnC^21^. Following ABT-263 treatment, Bcl-2 expression decreased, consistent with its mechanism of action. This decrease in Bcl-2 was accompanied by increased levels of annexin and caspases, indicating enhanced apoptosis-related signaling. The impact on mitochondrial function was associated with the negative regulation of oxidative phosphorylation and mitochondrial bioenergetics. Lagadinou et al. found that ABT-263 causes severe impairment of oxidative phosphorylation in acute myelogenous cells within minutes, leading to targeted apoptosis^22^. Interestingly, sensitivity to ABT-263 has been shown to be reactive oxygen species (ROS) dependent, with some cancer cells exhibiting enhanced sensitivity to ABT-263 due to higher intracellular ROS ^23^. As both aged skin cells and chronic SnCs share elevated intracellular ROS^24,25^, ABT-263 may be an effective senolytic for targeting skin SnCs to enhance regenerative capacity.

We observed a decrease in general senescence markers—p16, p21, and SA-β-gal—in the skin of aged mice after ABT-263 treatment. However, there were no changes in p16 or p21 gene expression in the skin of young 2-month-old mice, presumably because they have not accumulated a significant amount of SnCs. Despite the age-related reduction in senescence markers, ABT-263 treatment induced an increase in inflammation, characterized by significant macrophage infiltration of the treated skin. This heightened inflammation was supported by bulk RNA sequencing data, which revealed elevated gene expression in multiple inflammatory pathways. Given the overlap between many known SASP and general pro-inflammatory factors, it was not surprising that the transcription of some of these SASP genes (e.g., *Il1a, Il1b, Ccl3, Ccl8, Cxcl1, Cxcl2, Serpine1*) was significantly elevated following ABT-263 treatment. However, not all SASP factors were elevated; moreover, the murine SASP gene set in the Reactome pathway database was not found to be coordinately upregulated by Gene Set Enrichment Analysis (GSEA), which is in agreement with a finding by Saul et al ^26^ that there was no coordinate regulation of the human Reactome SASP gene set with respect to age in two separate cohorts.

Topical ABT-263 significantly altered gene expression in aged skin, particularly in pathways related to wound healing, including hemostasis, inflammation, angiogenesis, proliferation, and collagen and extracellular matrix regulation. This “priming” effect led to a notable improvement in wound healing kinetics, reducing the time to achieve complete healing. Accelerated healing was observable as early as day 6 and became statistically significant by day 15. Given the increased cellular infiltration, particularly with macrophages, after ABT-263 treatment, we hypothesize that this early inflammatory recruitment accelerates the wound healing process in aged mice. This macrophage recruitment might result from a senolysis-induced inflammatory response, as dying SnCs release their intracellular contents, which act as damage-associated molecular patterns (DAMPs) that activate the immune system. The transient inflammation induced by ABT-263 senolysis can help promote tissue repair and regeneration, as sensitization to inflammation has been shown to enhance wound healing, causing the skin to react faster when faced with a secondary injury^27^. Future work will delineate the origin and identities of the macrophages recruited after ABT-263-induced senolysis as they appear to be beneficial to aged wound healing.

Our results differ somewhat from other studies using ABT-263, which we believe is likely due to differences in the age of mice, delivery method (systemic versus topical), and time points examined after treatment. For instance, Kim et al. administered 5 μM ABT-263 dissolved in DMSO versus DMSO alone for five days via intradermal injections of in 25-month-old mice, which were sacrificed seven days post-injection^12^. They observed decreased levels of p16, p21, and SA-β-gal, but only certain SASP factors (*Il6, Mmp1, Cxcl1*) were significantly decreased, while others (*Mmp3, Mmp9, Mmp13, Tnf, Mcp1, Cxcl2*) showed no significant difference. They also noted a significant increase in Col1a1 expression, correlating with increased dermal thickness. Lagares et al. used 100mg/kg ABT-263 via 14-day oral gavage in a bleomycin dermal fibrosis mouse model^14^. Using FACS analysis of digested skin, they found an increase (though not statistically significant) in total leukocytes and inflammatory monocytes in ABT-263 treated young control (non-bleomycin treated) mice. Takaya et al. used a chimeric model with aged human skin on nude mice. They dosed ABT-263 intraperitoneally at 50 mg/kg/day in DMSO vs DMSO alone for 7 days per cycle for two cycles at 2-week intervals. Five weeks later, the graft was retrieved and demonstrated decreased p16, p53, and SA-β-gal. Utilizing a nude mouse model, this study excluded a normal murine inflammatory response and found decreased expression of SASP factors (*MMP3, MMP9, IL1α, IL6, TNFα*), but an increase in *Col1a1* expression and collagen density^17^. Overall, our study observed similar responses of aged skin to ABT-263, including reduced general senescence markers and increased collagen expression. However, the mixed acute inflammatory and SASP response was likely due to our brief five-day topical treatment and the early measurement of these markers.

The consistent increase in collagen expression in aged skin following ABT-263 treatment is intriguing. Aged skin generally demonstrates decreased collagen content (up to 75%) and increased fragmentation and disorganization ^28^. ABT-263 appears to enhance collagen expression and production, potentially improving aged skin structure and function. The removal of SnCs by ABT-263 may create a more favorable environment for healthy fibroblasts to produce collagen, thereby promoting tissue regeneration and repair. This observation contrasts with the effects of ABT-263 on fibrotic skin conditions, such as scleroderma and hypertrophic scars, where the treatment typically reduces collagen expression and targets αSMA-positive myofibroblasts ^14,18^. This difference in collagen response might be due to the distinct cellular environments and pathological processes involved in aging versus fibrotic conditions. In fibrotic skin, where excessive collagen deposition by myofibroblasts is pathologic, ABT-263 effectively reduces collagen levels, alleviating fibrosis. These contrasting outcomes highlight the context-dependent effects of ABT-263 and suggest its potential for selectively modulating collagen expression based on the specific skin condition being treated.

Our study underscores the potential of topical senolytic treatments to enhance wound healing in aging skin, presenting a potentially promising strategy for preoperative care. Pre-treating aged tissues with senolytics could potentially “prime” them for improved surgical wound healing in elective cases. By clearing chronic SnCs before surgery, senolytics may create a more favorable cellular environment for healing. This priming effect could enhance the efficiency of surgical procedures, particularly in elderly patients who often experience delayed healing and complications ^1,29^. While our findings are preliminary, numerous studies have demonstrated the ability of senolytics to effectively target SnCs. However, confirming these effects may necessitate exploring multiple senolytics alongside ABT-263. By broadening our investigation to senolytics with enhanced specificity, we aim to optimize treatments for diverse age groups and wound types. This approach could lead to tailored senotherapeutic strategies that maximize benefits while minimizing potential drawbacks.

## Methods

### Animals

All procedures were carried out under the National Institutes of Health’s Guide for the Care and Use of Laboratory Animals and approved by the Boston University Subcommittee on Research and Animal Care (202000006). Wild type aged (24 months) and young (2 month) C57BL/6 male mice were obtained from the National Institute of Health – National Institute of Aging.

### Topical senolytic treatment

Aged (24 months) mice were anesthetized with isoflurane anesthesia before topical senolytic treatment. Dorsal hair was removed with clippers followed by application of Nair™ (Church & Dwight Co., Inc, Ewing, NJ) hair removal lotion. 70% ethanol was used to cleanse the skin. Topical senolytic ABT-263 (5μM) or control vehicle-alone (DMSO) was applied topically on the skin and covered with a semi-occlusive dressing (3M™ Tegaderm™) for 5 days before wounding.

### Animal surgery

5 days after topical senolytic treatment, a 1 cm^2^ full-thickness excisional wound was created on the dorsal skin of mice under isoflurane anesthesia. 3M™ Tegaderm™ Transparent Film Dressing was used to cover the wounds. Mice were given 0.03 mg/kg buprenorphine subcutaneously for analgesia and were housed with food and water ad libitum. Postoperatively, wounds were imaged digitally every three days. ImageJ software was used to calculate the wound area (Version 1.53k).

### Real-time qPCR

The gentleMACS dissociator (Miltenyl Biotec) was used to homogenize the tissue and TRIzol™ Plus RNA Purification Kit (Catalog number: 12183555) was used to extract RNA from skin tissue isolated from aged male mice treated with ABT-263 and DMSO 5 days after treatment. RNA was reverse transcribed to cDNA using the Verso cDNA synthesis kit (ThermoScientific, ref. #AB-1453/B), following Nanodrop quantification. For quantitative real-time PCR, Maxima SYBR Green/ROX qPCR Master Mix (ThermoFisher Scientific, ref. #K0222) (Applied Biosystems) was used. To calculate the relative mRNA levels of senescence and SASP markers, β -actin was used as a control.

### Senescence-associated beta-galactosidase (SA-β-gal) stainin

5 days after ABT-263 and DMSO treatment. Tissues were first fixed with neutralized buffer formalin for 4-6 hours before being incubated overnight at 4ºC in a 30% sucrose solution. To detect SA-β-gal in skin tissue, the CellEvent™ Senescence Green Detection Kit (Invitrogen, cat. #C10851) was used. Frozen 5-μm-thick sections were washed with buffer and then with 1% BSA. Sections were treated with a prewarmed working solution and placed in a humidified chamber to prevent moisture loss before being incubated overnight at 37ºC without CO_2_ and protected from light. After washing in buffer, the sections were mounted with VECTASHIELD® Antifade Mounting Medium with DAPI (H-1200-10). Nikon Eclipse E400 fluorescent microscope at 10X and 20X magnification was used to capture microscopic images. The SA-β-gal positive cells were counted using Fiji-ImageJ (Version 1.53c).

### Immunohistochemistry and Histological Assessment

After fixation in 4% paraformaldehyde in PBS for 2 h at 4°C with subsequent overnight sucrose treatment at 4°C, the 5-μm-thick cryosections were washed in PBS. Antigen retrieval was performed in EnVision FLEX Target Retrieval Solution Low pH (Dako, ref. #K8005) using Dako PT Link antigen retrieval machine. Sections were washed with PBS and traced with a hydrophobic pen. For immunohistochemistry, sections were incubated in BLOXALL^®^ Blocking Solution (SP-6000) for 10 mins for the quenching of endogenous peroxidase activity. Blocking was performed for 20 mins at room temperature with 2.5% horse serum. Sections were incubated with rabbit anti-p21 primary antibodies (ab188224); CD4 (ab183685), CD8 (CST 98941), F4/80 (ab6640) primary antibodies overnight at 4°C (1:100), washed with PBS, followed by incubation with ImmPRESS Polymer Reagent provided in the ImmPRESS^®^ Horse Anti-Rabbit IgG Polymer Kit (Vector Laboratories: Cat. No.: MP-7401) for 30 mins. After aspiration of secondary antibodies and extensive washing to remove unbound antibodies, sections were incubated with peroxidase (HRP) detection system (ImmPACT® DAB Peroxidase Substrate, Vector Laboratories: Cat. No.: SK-4105) for 30 seconds to 1 minute or until desired stain intensity develops. The sections were rinsed in tap water and counterstained with Hematoxylin, mounted with VECTASHIELD® Antifade Mounting Medium (H-1000-10), and coverslipped. Brightfield microscopy images were captured using Nikon Eclipse E400 fluorescent microscope at 10x SA-β-gal and anti-p21 staining, respectively. Percentages of SA-β-gal- and p21-positive cells were calculated using ImageJ software.

For Hematoxylin and Eosin staining, deparaffinized sections were rehydrated with distilled water. Mayers Hematoxylin staining for 1 minute Wash with distilled water, then counterstain with Alcoholic-Eosin for 1 minute before dehydrating with Ethanol. After that, the stained slides were mounted and coverslipped. A Nikon Eclipse E400 fluorescent microscope was used to capture brightfield microscopy images at a magnification of 10x.

### Bulk RNA Sequencing and Analysis

FASTQ files were aligned to mouse genome build mm10 using STAR(version 2.7.9a)^30^. Ensembl-Gene-level counts for non-mitochondrial genes were generated using featureCounts (Subread package, version 1.6.2) and Ensembl annotation build 109 (uniquely aligned proper pairs, same strand). FASTQ quality was assessed using FastQC (version 0.11.7), and alignment quality was assessed using RSeQC (version 3.0.0). Variance-stabilizing transformation (VST) was accomplished using the varianceStabilizingTransformation function in the DESeq2 R package (version 1.23.10)^31^. Differential expression was assessed using the Wald test implemented in the DESeq2 R package. Correction for multiple hypothesis testing was accomplished using the Benjamini-Hochberg false discovery rate (FDR). All analyses were performed using the R environment for statistical computing (version 4.1.2).

Gene Set Enrichment Analysis (GSEA) (version 2.2.1)^32^ was used to identify biological terms, pathways and processes that are coordinately up- or down-regulated within each pairwise comparison. The Entrez Gene identifiers of all genes in the Ensembl Gene annotation were ranked by the Wald statistic computed between the DMSO and ABT-263 groups. Ensembl Genes matching multiple mouse Entrez Gene identifiers were excluded prior to ranking. This ranked list was then used to perform pre-ranked GSEA analyses (default parameters with random seed 1234) using the Entrez Gene versions of the MH (Hallmark), MH CP (Biocarta, Reactome, WikiPathways), and M5 (Gene Ontology, GO) gene sets obtained from the Molecular Signatures Database (MSigDB), version 2023.2.Mm.

## Statistical Analysis

Unpaired t-test was used for comparisons between ABT-263 and DMSO-treated animals. The results are presented as mean + SEM, with p<0.05 considered statistically significant.

## Data availability

The data discussed in this publication have been deposited in NCBI’s Gene Expression Omnibus and are accessible through GEO Series accession number GSE271149 (https://www.ncbi.nlm.nih.gov/geo/query/acc.cgi?acc=GSE271149). Data supporting the findings of this study are available from the corresponding author upon reasonable request.

**There are no conflicts of interests to declare for any authors**

## Acknowledgements

We would like to thank the BU Microarray and Sequencing Resource for assistance in experimental design and bulk RNA sequencing support.

## Supplementary data

**Supplementary Figure 1. Topical ABT-263 does not alter p16 or p21 expression in young mouse skin**. A) p16 and p21 gene expression relative to β-actin after 5d of ABT-263 (N=5) vs DMSO (N=5), 2 month old mice. t-test, * p<0.05.

**Supplementary Figure 2. Topical ABT-263 decreased CD4 and CD8 cell infiltration in aged skin**. CD4 and CD8 staining of skin after 5d of ABT-263 (N=5) vs DMSO (N=5), 24 month old mice. Number of cells/high powered field. t-test, * p<0.05 significance level.

**Supplementary Figure 3. Topical ABT-263 effects on systemic blood counts in aged mice**. Blood was obtained one day after 5 days of topical treatment with ABT-263 (N=3) vs DMSO (N=2) in 24 month old mice. T test, * p<0.05 significance level.

**Supplementary Table 1. Gene-level analysis results**. This Excel file contains gene annotation (columns A-E), fold changes and Wald test results (columns F-I), counts of uniquely aligned read pairs assigned to each gene (columns J-P), and variance-stabilizing-transformed (VST) expression values (columns Q-W). Counts and VST values are shaded according to VST expression values, where blue and red indicate VST values that are ≥ 2 standard deviations below or above the mean (white) of each row, respectively.

**Supplementary Table 2. GSEA results**. This Excel file contains the MSigDB collection (column A), name and size (columns B-C), Normalized Enrichment Score (NES, column D), and nominal p and FDR q values (columns E-F) for each gene set (version 2023.2.Mm).

