## supplemental figures for "Topical ABT-263 treatment reduces aged skin senescence and improves subsequent wound healing"

#### SUPPLEMENTAL FIGURE 1

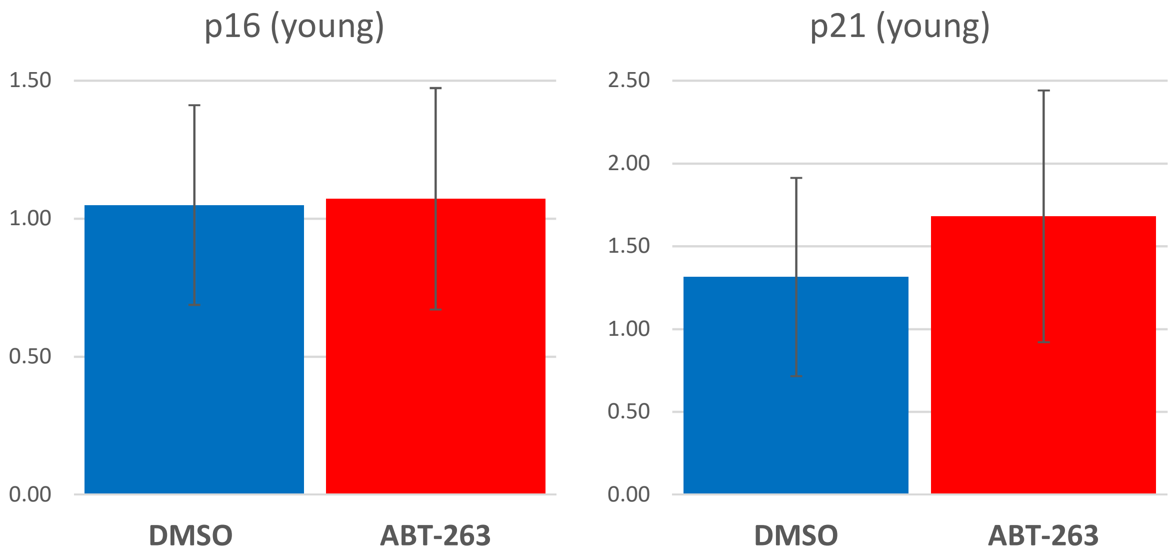


**Supplementary Figure 1. Topical ABT-263 does not alter p16 or p21 expression in young mouse skin.** A) p16 and p21 gene expression relative to β-actin after 5d of ABT-263 (N=5) vs DMSO (N=5), 2 month old mice. t-test, * $p<0.05$.

#### SUPPLEMENTAL FIGURE 2

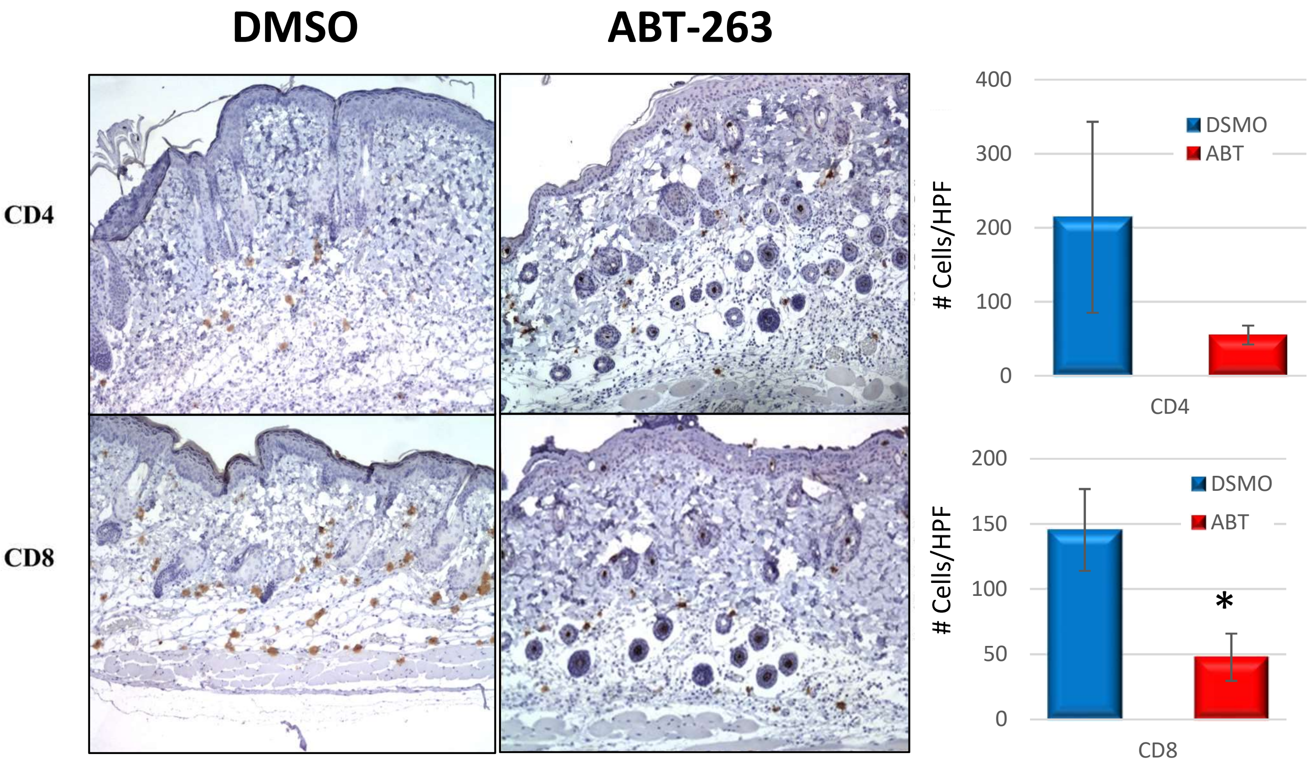


**Supplementary Figure 2. Topical ABT-263 decreased CD4 and CD8 cell infiltration in aged skin.** CD4 and CD8 staining of skin after 5d of ABT-263 (N=5) vs DMSO (N=5), 24 month old mice. Number of cells/high-powered field. t-test, * $p<0.05$ significance level.

### SUPPLEMENTAL FIGURE 3

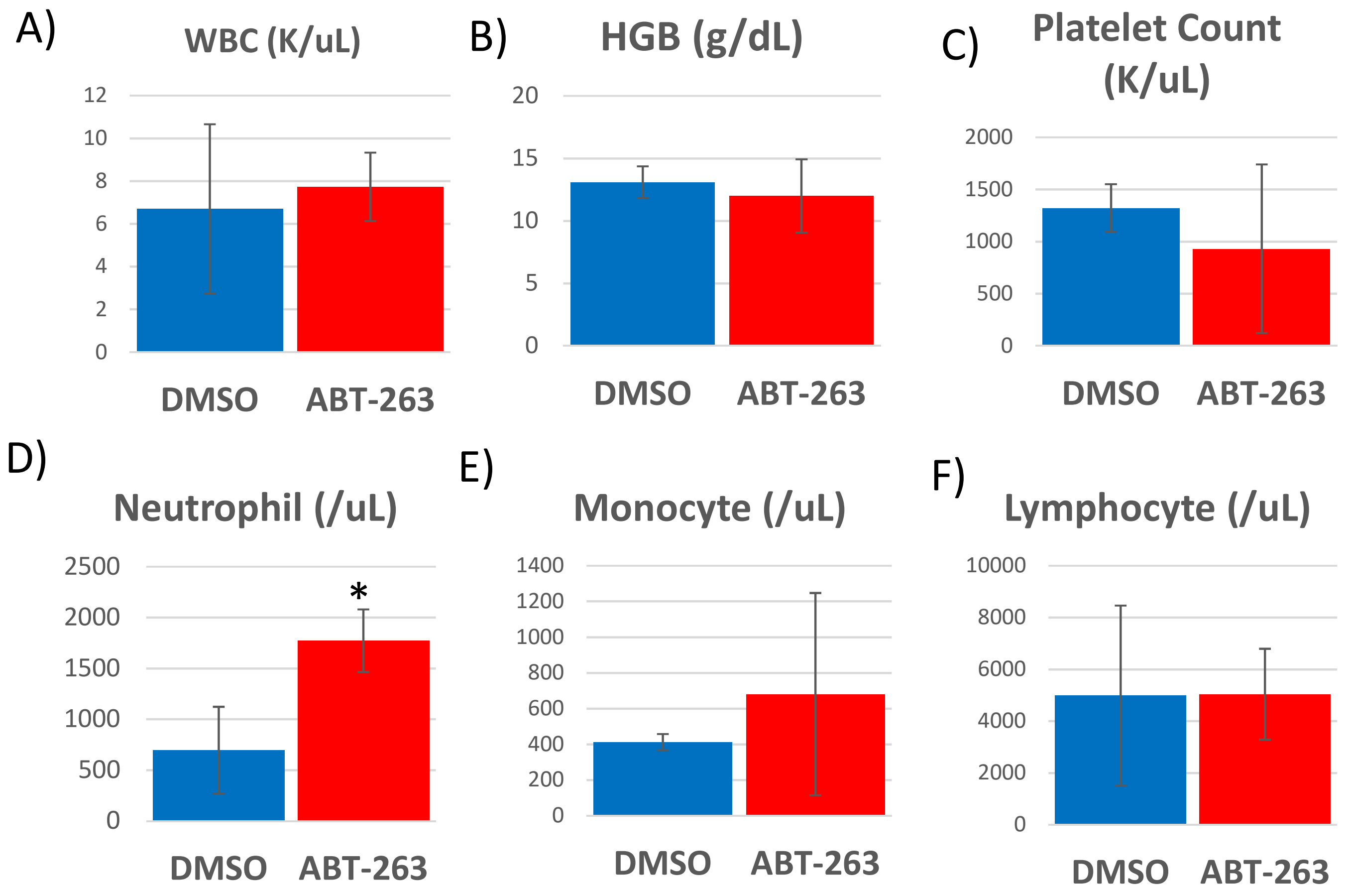


**Supplementary Figure 3. Topical ABT-263 effects on systemic blood counts in aged mice.** Blood was obtained one day after 5 days of topical treatment with ABT-263 (N=3) vs DMSO (N=2) in 24 month old mice. t-test, * $p<0.05$ significance level.
